# CD38 is a good predictor of anti-PD-1 immunotherapy responsiveness in hepatocellular carcinoma

**DOI:** 10.1101/638981

**Authors:** Siting Goh, Harry Ho Man Ng, Valerie Chew, Xin Ni Sim, Huihua Li, Sherlly Lim, Jeffrey Chun Tatt Lim, Josh Jie Hua Loh, Khin Sabai, Clara Chong Hui Ong, Tracy Loh, Wei Qiang Leow, Joycelyn Lee Jie Xin, Han Chong Toh, Fabio Malavasi, David Wai Meng Tai, Ser Yee Lee, Pierce Chow, Evan Newell, Su Pin Choo, Joe Yeong, Tony Kiat Hon Lim

## Abstract

Hepatocellular carcinoma (HCC) is the fourth leading cause of cancer-associated mortality in the world. However, with the associated low five-year survival and high recurrence rates, alternative treatment modalities specifically immunotherapy have been researched. A correlation between CD38^+^ tumour-infiltrating leukocyte (TIL) density and improved prognosis was found in a recent study. However, studies relating to CD38 expression in immune infiltrates within tumours are limited. In the present study, we confirmed the expression of CD38 on macrophages in HCC and determined the relationship between CD38^+^ leukocytes and lymphocytes and patient response to immunotherapy. Using immunohistochemistry, we analysed tissue samples obtained from 20 patients from Singapore with HCC prior to immunotherapy. Tumour infiltrating leukocytes expression within tumour were correlated to the responsiveness of patients to immunotherapy.

Expression of CD38 was found within the tumour cells and surrounding immune infiltrates including lymphocytes and macrophages. We then ask whether CD38 expression by the distinct cell populations may acquire theranostic relevance. Patients with higher level of CD38^+^ immune infiltrate subsets had significantly better response to anti-PD-1 immunotherapy, and this is also true for CD38^+^ lymphocytes within the tumour microenvironment. In particular, a cut-off of 13.0% positive out of total leukocytes and 12.4% positive out of total lymphocytes is found to be of strong predictive value of responsiveness to immunotherapy treatment, thus a strong theranostic impact is seen by using CD38 as a biomarker for anti-PD-1 therapy.

The establishment of an association between CD38 expression and the response to anti-PD-1 immunotherapy in HCC, could be applied to a larger cohort outside Singapore. These may eventually change the routine testing in clinical practice to identify HCC patients suitable for immunotherapy.

## 1 Introduction

Hepatocellular carcinoma (HCC) is the fifth and ninth most frequently diagnosed cancer in adult males and females, respectively,^1^ and is ranked as the fourth leading cause of cancer-associated mortality in the world. Cirrhosis is a major risk factor for HCC, and this is often caused by chronic hepatitis B or C infection. Surgical resection and liver transplantation are curative therapeutic options for early-stage HCC. However, five-year survival rates following surgical resection for early stage disease remain relatively low (17%-53%), with a recurrence rate as high as 70%.^2–4^

Currently, the first-line therapy for advanced HCC is oral Sorafenib and Lenvantinib^5, 6^. They are both oral multi-kinase inhibitor (MKI), capitalizing on vasoendothelial growth factor (VEGF) inhibition for HCC. Although, these have largely prolonged the survival of patients, a large proportion of patients diagnosed with intermediate and advanced stage HCC still reported a median overall survival of 21.1 months^6^ and 10.7 months^7^ respectively in large randomized trials. Thereby highlighting the necessity of other options to treat refractive advanced HCC patients.

Besides VEGF inhibition with oral MKI, PD-1/PDL-1 and CTLA-4 inhibition has emerged as an exciting therapeutic strategy in systemic armamentarium for HCC. Immunotherapy utilising checkpoint blockade antibodies has delivered promising results in diseases such as melanoma^8^ and lung cancer^9^. This has led to FDA approval for treatments using monoclonal antibodies, including nivolumab, ipilimumab, pembrolizumab, and atezolizumab, against specific checkpoint molecules, such as cytotoxic T-lymphocyte–associated antigen 4 (CTLA-4), programmed cell death protein-1 (PD-1) and programmed death-ligand 1 (PD-L1).

The efficacy and safety of nivolumab, which targets PD-1, have been explored in a phase I/II trial (CheckMate 040). The preliminary results of this trial were promising, and that led to the FDA granting accelerated approval of nivolumab treatment for HCC patients previously treated with sorafenib^10, 11^.

Current data showed phase I/II trials with response rates about 20%, and thus a number of different biomarkers that have been proposed to differentiate patients who will benefit from PD-1 immunotherapy. In addition to PD-L1 expression as a biomarker for response to immunotherapy,^12–14^ tumour mutation burden (TMB) and microsatellite instability (MSI) have also been proposed as predictive markers for immunotherapy, with these markers depicting the number of neoantigens in the tumour that would potentially be recognized by the immune system.^15^ IFNγ gene signature is also used to potentially discriminate responsiveness to PD-1 checkpoint blockade.^16–19^ However, in HCC the use of PDL-1 – cut-off 1% on tumour cells has its limitations in predicting response.^10^ Thus, there remains to be seen if there is any robust biomarker in HCC.

The tumour microenvironment (TME) has also become of interest to the field of immunotherapy. Under normal conditions, tissue homeostasis acts as barrier against tumour formation, with tumours altering the stromal components during their development and metastasis. The tumour microenvironment is hypoxic and immunosuppressive, and a multifunctional molecule called CD38 is involved in this mechanism. CD38 structurally resembles CD1a and serves as an ectozyme in the adenosinergic pathway.^20^ In hypoxic environments, NAD^+^ is released by the salvage pathway and hydrolysed by CD38 to form adenosine diphosphate ribose. This is further degraded to adenosine monophosphate (AMP) through the CD38-CD203a-CD73 pathway. Following this, CD73 dephosphorylates AMP to adenosine.^21, 22^ Accumulated extracellular adenosine then binds to various receptors on a range of immune cells, impeding their infiltration and activation.^23, 24^ This represents an alternative immunosuppressive mechanism to the PD-1/PD-L1 pathway. Inhibition of the adenosine pathway has been shown to weaken the intensity of immunosuppression in the TME.^25^ Furthermore, reversal of hypoxia via oxygen supplementation in a murine model resulted in a significant reduction in solid tumour growth and metastasis.^26^ Similarly, co-treatment with PD-1 blockade and adenosine receptor inhibitors has been found to improve immune cell responses and result in increased tumour suppression in various mouse models.^27–29^

Beside serving as a ectozyme, CD38 also functions as a surface membrane marker in various immune cells and non-lymphoid tissues.^30^ The relevance of CD38 to HCC was established in a recent study by our group, where a correlation between CD38^+^ tumour-infiltrating leukocyte (TIL) density and improved prognosis was found.^31^ The expression of CD38 has been reported in a range of immune cell populations,^30^ but data regarding macrophages and lymphocytes are limited.

In the present study, we confirmed the expression of CD38 on immune cells and tumours in HCC. Following this, we determined the relationship between CD38^+^ leukocytes and lymphocytes and patient response to immunotherapy in a pilot, retrospective cohort of Asian HCC patients (n=20).

## 2. Materials and Methods

### 2.1 Patients and tumours

These were a total of 20 archival formalin-fixed, paraffin-embedded (FFPE) specimens taken from Asian patients with HCC undergoing immunotherapy between January 2015 and December 2018 at the Department of Medical Oncology, National Cancer Centre. Samples were all obtained prior to immunotherapy at the Department of Anatomical Pathology, Division of Pathology, Singapore General Hospital. Surgical resected samples or biopsy specimens were obtained prior to PD-1/PD-L1 therapy. Tumours were staged and graded according to the BCLC staging^32^ or AJCC staging system^33^ and the Edmonson-Steiner grading system.^34^ The Centralized Institutional Review Board of SingHealth provided ethical approval for the use of patient materials in this study (CIRB ref: 2014/590/B).

### 2.2 Immunohistochemistry (IHC)

IHC was performed on the FFPE tissue samples as previously described.^35^ TMA sections of 4μm thickness were incubated with antibodies specific for CD8, CD38, CD68 and cell nucleus, as listed in Supplementary Table 1. Appropriate positive and negative controls were included. To generate the scoring of antibody-labeled sections, images were captured using an IntelliSite Ultra-Fast Scanner (Philips, Eindhoven, Netherlands). The percentage of leukocyte cells displaying unequivocal staining of any intensity for CD38 was determined by pathologists blinded to clinicopathological and survival information.

### 2.3 mIF analysis of TMAs

Multiplex immunofluorescence/immunohistochemistry (mIF or mIHC) was performed using an Opal Multiplex fIHC kit (PerkinElmer, Inc., Waltham, MA, USA), as previously described by our group and other studies.^31, 36–47^ TMA sections (4 µm thick) were labelled with primary antibodies against CD38, CD8 and CD68, followed by appropriate secondary antibodies. All antibodies used are listed in Supplementary Table 1. This was followed by the application of a fluorophore-conjugated tyramide signal amplification buffer (PerkinElmer, Inc.) and the nuclear counterstain DAPI. A Vectra 3 pathology imaging system microscope (PerkinElmer, Inc.) was used to obtain images, and these were analysed using inForm software (version 2.4.2; PerkinElmer, Inc.).^48–51^

### 2.4 Validation, follow-up and statistical analysis

Follow-up data were obtained from medical records. An unpaired, two-tailed Student’s t-test was used to compare CD38 expression in patients with that was successfully treated by immunotherapy and patients whom were not. The associations between clinicopathological parameters and the frequency of CD38^+^ leukocytes were analysed using χ² and Fisher’s exact tests. Statistical analysis was performed using RStudio 1.1.456 running R 3.5.0^52, 53^ (R-core Team, R Foundation for Statistical Computing, Vienna, Austria) and GraphPad Prism 8.0.0 for Windows (GraphPad Software, Inc., San Diego, CA, USA). P<0.05 was considered to indicate a statistically significant difference.

## 3. Results

### 3.1 Tumour, lymphocytes and macrophages express CD38, and macrophages represent the largest CD38^+^ immune cell subset in HCC

We initially sought to verify the expression of CD38 by different immune infiltrates in the HCC samples. A high level of co-localisation between CD38 and CD68 expression was visualised under mIF/mIHC (Fig. 1A), indicating that CD38 is expressed in macrophages. CD38 was also expressed in CD8^+^ lymphocytes (Fig. 1B), and in tumour cells (Fig. 1C). CD38 staining is seen on tumour cells for 2 out of 36 patients in IHC staining. DAPI Nucleus staining was used to identify the cells. Further analysis showed that expression of CD38^+^ immune infiltrates in HCC were more in non-responders when compared to those who responded to anti-PD-1/PD-L1 treatment (Fig. 2).

**Figure 1.**
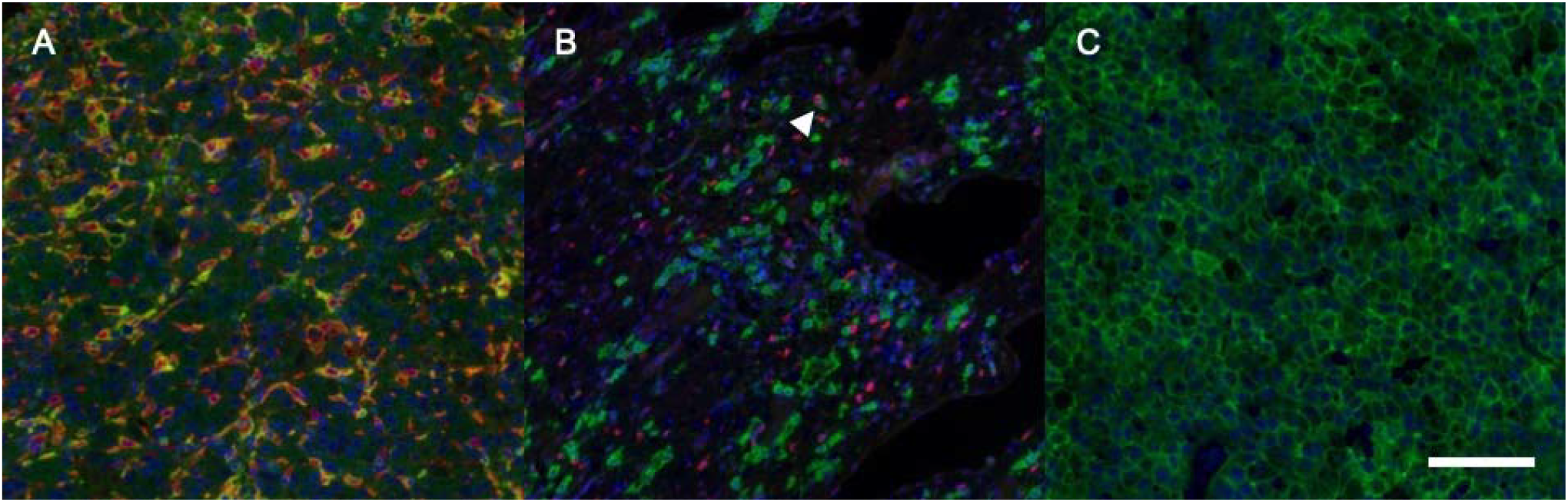
Expression of CD38 in different cell types within formalin-fixed, paraffin-embedded HCC sections. (A) Multiplex immunohistochemistry revealed that CD38 (green) is frequently co-localized with the macrophage marker CD68 (red) in the HCC tumour microenvironment. (B) Similar co-localization (white arrow) between CD38 (green) and CD8^+^ lymphocytes (red) was also observed. (C) CD38 (green) is also expressed in HCC tumour cells. HCC, hepatocellular carcinoma. DAPI staining for cell nuclei (blue)

**Figure 2.**
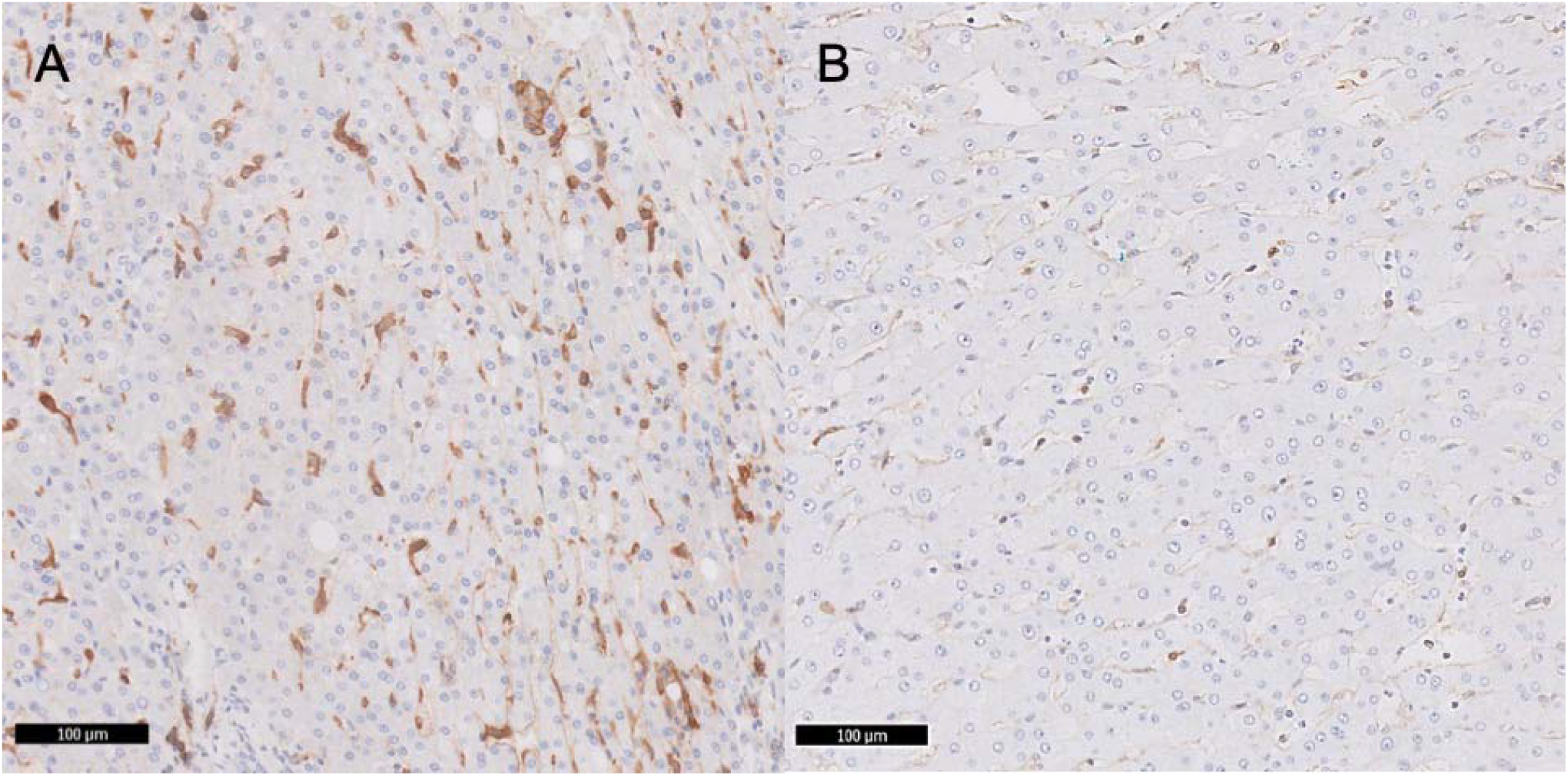
The responders harboured more CD38^+^ cells, compared to the non-responders of PD-1 treated HCC. Respective CD38 immunohistochemistry staining showing (A) responders harboured more CD38^+^ cells compared to the (B) non-responders.

### 3.2 CD38+ immune infiltrates are associated with partial responsiveness to anti-PD-1 immunotherapy treatment in HCC

We further assessed whether the presence of CD38^+^ immune infiltrate subsets in specific locations affects the responsiveness of patients with HCC to nivolumab. In our study, the response criteria were classified as either response present (including complete and partial response) or no response (including both stable and progressive disease). The response criteria are determined according to the RECIST 1.1 guidelines^54^ with radiology review. The presence of CD38^+^ immune infiltrate subsets within the tumour was strongly associated with a positive response to anti-PD-1 immunotherapy (responsive vs. non-responsive, 15.83±5.943% vs. 3.539±1.070%; p=0.0041; Fig. 3). Similarly, CD38^+^ lymphocytes (responsive vs. non-responsive, 26.13±10.84% vs. 4.495±1.437%; p=0.0034) was also strongly associated with better responsiveness in anti-PD-1 therapy specifically, but not CD38^+^ macrophages (responsive vs. non-responsive, 5.536±2.521% vs. 2.582±1.049%; p=0.2160).

**Figure 3.**
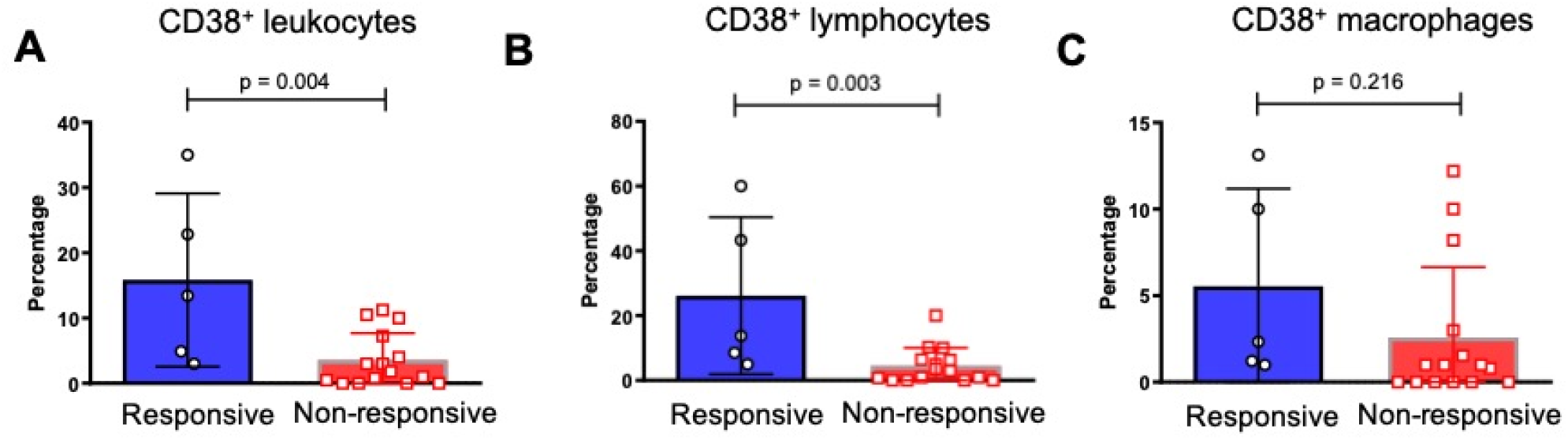
Therapeutic response of HCC patients in relation to the CD38^+^ immune infiltrate subsets. (A) The percentage of CD38^+^ immune infiltrate subsets within the tumour in responders and non-responders to anti-PD-1 immunotherapy. (B) The percentage of CD38^+^ lymphocytes within the tumour in responders and non-responders to anti-PD-1 immunotherapy. (C) The percentage of CD38^+^ macrophages within the tumour in responders and non-responders to anti-PD-1 immunotherapy. Data are presented as the mean ± standard error mean. (n = 20 formalin-fixed, paraffin-embedded samples).

### 3.3 CD38^+^ leukocyte and lymphocyte proportion is of strong predictive value of responsiveness to treatment

As shown in Fig. 4, the optimal cut-off for CD38^+^ leukocyte proportion has been defined using receiver operating characteristic analysis. The cut-off used was 4.4% positive out of total immune infiltrates, and this cut-off achieved 75.0% accuracy, 73.3% specificity and 80.0% sensitivity. The area under curve (AUC) is 0.867. Similarly, the optimal cut-off for CD38^+^ lymphocytes proportion was defined. The cut-off used was 7.5% positive out of total lymphocytes, and this cut-off was achieved 80.0% accuracy, 80.0% specificity and 80.0% sensitivity.

**Figure 4.**
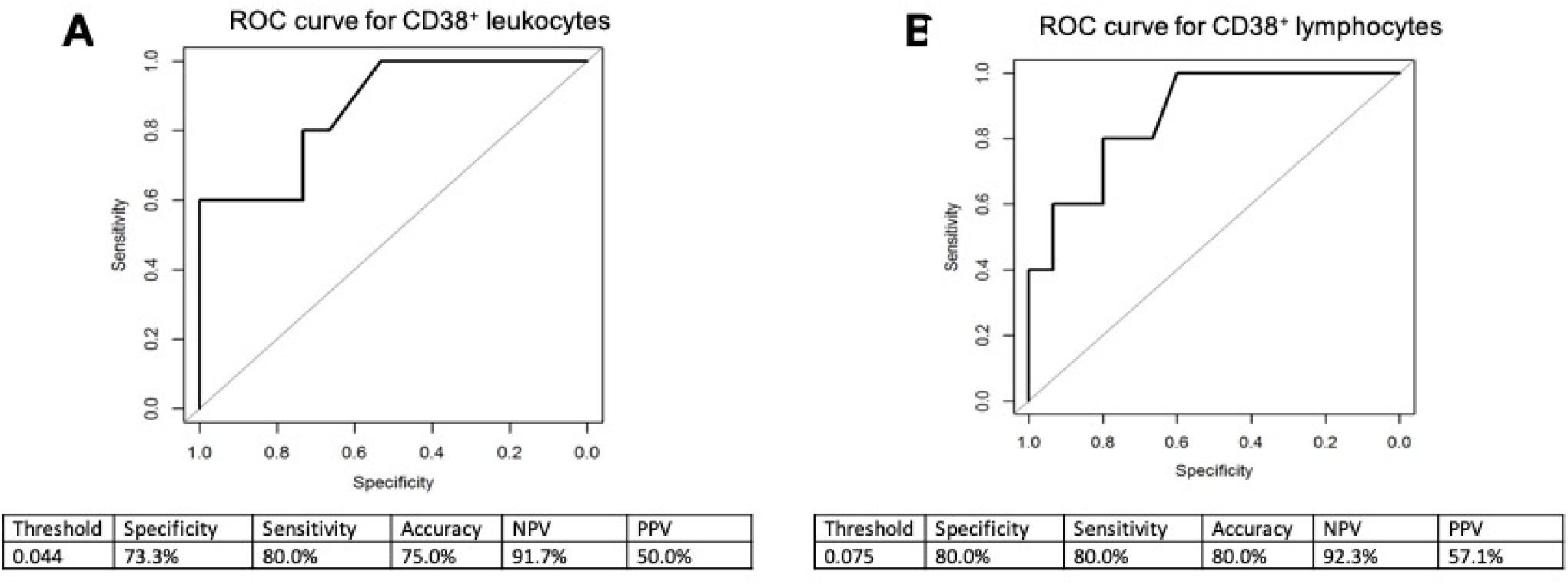
Receiver operating characteristic curve for the ability of CD38^+^ leukocyte and lymphocyte proportion to identify responders. (A) CD38^+^ leukocyte: AUC=0.867 (0.679, 1.000), (B) CD38^+^ lymphocytes: AUC=0.873 (0.704, 1.000). Sensitivity refers to the proportion of true positive subjects with the disease among subjects with disease. Specificity refers to the proportion of true negative subjects without the disease among subjects without disease. PPV refers to the proportion of patients with positive results among subjects with positive results. NPV refers to the proportion of subjects without disease with a negative result among subjects with negative results. Accuracy refers to the proportion of subjects correctly classified among all subjects. AUC, area under the curve; PPV, positive predictive value, NPV, negative predictive value.

## 4. Discussion

Cancer immunotherapy is mechanistically different to other treatment modalities, such as cytotoxic therapies and small module inhibitors, as it targets the TME rather than the tumour itself. So far, minimal side effects have been identified, and the potential for application to different types of cancer seems particularly promising. However, the overall patient response rate to PD-1/PD-L1 inhibitors remains unsatisfactory, limiting its application in clinical practice. This phenomenon may be the result of variability in the immune microenvironment between different cancers. Thus, further investigation of biomarkers is of the utmost importance to fully understand any associations with clinical outcomes, and identify patients most likely to benefit from treatment.

Multiple studies have demonstrated correlations between therapeutic response rates and PD-L1 expression in tumours, which are likely due to the close relationship between PD-L1 and PD-1. Increased PD-L1 expression is generally believed to be associated with increased response rate and improved clinical benefit in PD-1 blockade therapy. However, the conclusions drawn from multiple trials have not always been consistent.^55–60^ Thus, investigation of alternative immunosuppression pathways to PD-1/CTLA-4 is necessary. One such immunosuppressive mechanism proposed to be of relevance is the adenosinergic pathway, where extracellular adenosine exerts local immunosuppression through tumour-intrinsic and host-mediated mechanisms.

The adenosinergic pathway involves CD38, which is a multifunctional marker that is expressed in various regulatory cells, including myeloid-derived suppressor cells, mesenchymal stem cells and NK cells.^61^ In a recent study, CD38 was found to be expressed by a subset of tumours with high levels of basal or treatment-induced infiltration.^62^ In the present study, multiplex IHC revealed the expression of CD38 on the surface of both leukocytes and HCC tumour cells (Fig. 1), which is consistent with the current literature.^30, 62^ Further previous studies have demonstrated that tumours treated with PD-1/PDL-1-specific antibodies develop treatment resistance through upregulation of CD38, which follows the release of all-trans retinoic acid and IFNβ in the TME. This results in the suppression of CD8^+^ T cell function via the adenosine signalling pathway.^62^ The role of adenosine in immune exhaustion and the observed expression of CD38 on immune infiltrates in the present study suggest the presence of a complex interplay between the inflammatory response and immune suppression via adenosine production. Thus, this immunosuppression mechanism may represent a promising target for immunotherapy.

Notably, the expression of CD38 was observed on macrophages in human HCC tumour samples. Previously, CD38 was found to be in macrophages isolated from mice^63, 64^, cell lines^65^ and on human *ex vivo* experiments^66^, but there is not much direct evidence on CD38 expression on macrophages in humans. This study confirmed that CD38 is expressed on tumour cells as well as multiple types of immune cells, including macrophages. Further analysis on CD38 expression established that responsiveness to immunotherapy is associated with higher levels of CD38^+^ immune infiltrates within the TME. This is also true for CD38^+^ lymphocytes levels within the microenvironment but the CD38^+^ macrophages subset did not achieve significance (Fig. 3), suggesting a role of CD38^+^ lymphocytes in affecting the response of immunotherapy. CD38 is shown to play an important role in lymphocyte activation.^67^ Our lab’s previous studies have ascertained the role of activated lymphocytes and CD38 in HCC prognosis,^31^ and expression of CD38 in lymphocytes has been shown to be a marker in other cancers.^68^ Considering the role of CD38 in the adenosine signaling pathway during hypoxia and the involvement of TILs in pro-inflammatory process, it is possible with anti-PD-1 immunotherapy, these CD38^+^ lymphocytes are suppressed thereby allowing favourable therapeutic responses.

In addition to the usage of PD-1-specific antibodies to treat HCC, other trials have also investigated whether combination immunotherapy can be used to overcome tumour resistance. One such trial is a phase Ib randomised clinical study, evaluating the safety and efficacy of administrating the PD-L1- specific antibody atezolizumab with bevacizumab, which is a monoclonal antibody that targets VEGF, as a treatment for HCC. This is an on-going trial that is due for completion in 2021.^69^ Potential future directions could include assessing the effects of CD38^+^ leukocyte density on the response to combined immunotherapy.

As previously stated, despite an abundance of biomarkers were proposed as a biomarker for immunotherapy therapy in other cancers, it is yet to been seen if there has been any biomarker for HCC. Our paper has successful demonstrated that CD38^+^ immune subsets or lymphocytes may also be useful as a biomarker to predict anti-PD-1 immunotherapy response. More studies are needed to confirm this phenomenon. Therefore, we propose to validate further by adopting CD38 IHC or mIF/mIHC staining in clinical practice to identify these patients who will gain benefits remarkably by this adjunctive test as the implementation of personalised medicine.

Limitations of the present study include a limited sample size which should be a common limitation for anti-PD-1 immunotherapy study in HCC, but the predictive significance between responders and non-responders were substantial. Moreover, a proportion of our patient cohort received PD-1 immunotherapy with another agent, making it a heterogenous population. However, this also reflects more closely to real life clinical practice, as most of the patients would receive combined therapy. Further studies may also be needed to investigate the effect of CD157 on response to anti-PD-1 immunotherapy, as this is a CD38 paralogue. The two molecules possess dual receptorial and NADase functions, and CD157 is widely expressed across lymphoid tissues, including immune cells such as lymphocytes and macrophages.^70^

In conclusion, the present study established an association between CD38 expression and the response to anti-PD-1 immunotherapy in HCC. Future investigations will look to apply this to a larger cohort or outside Singapore, and make comparisons with a non-Asian cohort. The eventual aim is to apply these findings as a routine test in clinical practice, to identify patients suitable for immunotherapy.

## 5. Declarations

### 5.1 Ethics approval and consent to participate

The Centralized Institutional Review Board of SingHealth provided ethical approval for the use of patient materials in this study (CIRB ref: 2014/590/B).

### 5.4 Author Contributions

JY and TL conceived, directed and supervised the study. HHMN and SG collated and interpreted the data and performed biostatistical analysis. XNS and JJHL constructed TMAs and performed IHC. HHMN and JY performed immunohistochemical scoring. FM, KS, CO, TL and WQL contributed to the scientific content of the study. DT, JLJX, SPC and HCT provided scientific inputs from Oncology perspectives. SYL and PC provided scientific inputs from Surgery perspectives. SG and HHMN drafted the manuscript with the assistance of JY, with final review from all authors.

## 5.5 Acknowledgements

We thank the funding bodies such as the Centre Grant of Singapore General Hospital (grant no. NMRC/CG/M011/2017_SGH, NMRC/CIRG/1454/2016) and the AM-ETHOS Duke-NUS Medical Student Fellowship Award (grant no. AM-ETHOS01/FY2018/10-A10). We also thank Dr. Alice Bridges, Dr. Lam Jianhang, Mr. Lim Chun Chye and Dr. Lim Tong Seng for critical review of the manuscript.

## References

1. Global Burden of Disease Liver Cancer C, Akinyemiju T, Abera S, et al. The Burden of Primary Liver Cancer and Underlying Etiologies From 1990 to 2015 at the Global, Regional, and National Level: Results From the Global Burden of Disease Study 2015. JAMA Oncol 2017;3:1683–1691.

2. Zheng J, Chou JF, Gonen M, et al. Prediction of Hepatocellular Carcinoma Recurrence Beyond Milan Criteria After Resection: Validation of a Clinical Risk Score in an International Cohort. Ann Surg 2017;266:693–701.

3. Chen XP, Qiu FZ, Wu ZD, et al. Long-term outcome of resection of large hepatocellular carcinoma. Br J Surg 2006;93:600–6.

4. Ruan DY, Lin ZX, Wang TT, et al. Nomogram for preoperative estimation of long-term survival of patients who underwent curative resection with hepatocellular carcinoma beyond Barcelona clinic liver cancer stage A1. Oncotarget 2016;7:61378–61389.

5. Kudo M, Finn RS, Qin S, et al. Lenvatinib versus sorafenib in first-line treatment of patients with unresectable hepatocellular carcinoma: a randomised phase 3 non-inferiority trial. Lancet 2018;391:1163–1173.

6. Llovet JM, Ricci S, Mazzaferro V, et al. Sorafenib in advanced hepatocellular carcinoma. N Engl J Med 2008;359:378–90.

7. Meyer T, Fox R, Ma YT, et al. Sorafenib in combination with transarterial chemoembolisation in patients with unresectable hepatocellular carcinoma (TACE 2): a randomised placebo-controlled, double-blind, phase 3 trial. Lancet Gastroenterol Hepatol 2017;2:565–575.

8. Zhang B, Zhou YL, Chen X, et al. Efficacy and safety of CTLA-4 inhibitors combined with PD-1 inhibitors or chemotherapy in patients with advanced melanoma. Int Immunopharmacol 2019;68:131–136.

9. Gamerith G, Kocher F, Rudzki J, et al. ASCO 2018 NSCLC highlights-combination therapy is key. Memo 2018;11:266–271.

10. El-Khoueiry AB, Sangro B, Yau T, et al. Nivolumab in patients with advanced hepatocellular carcinoma (CheckMate 040): an open-label, non-comparative, phase 1/2 dose escalation and expansion trial. Lancet 2017;389:2492–2502.

11. Zhu AX, Finn RS, Edeline J, et al. Pembrolizumab in patients with advanced hepatocellular carcinoma previously treated with sorafenib (KEYNOTE-224): a non-randomised, open-label phase 2 trial. Lancet Oncol 2018;19:940–952.

12. Grigg C, Rizvi NA. PD-L1 biomarker testing for non-small cell lung cancer: truth or fiction? J Immunother Cancer 2016;4:48.

13. Teixido C, Vilarino N, Reyes R, et al. PD-L1 expression testing in non-small cell lung cancer. Ther Adv Med Oncol 2018;10:1758835918763493.

14. Udall M, Rizzo M, Kenny J, et al. PD-L1 diagnostic tests: a systematic literature review of scoring algorithms and test-validation metrics. Diagn Pathol 2018;13:12.

15. Chang L, Chang M, Chang HM, et al. Microsatellite Instability: A Predictive Biomarker for Cancer Immunotherapy. Appl Immunohistochem Mol Morphol 2018;26:e15–e21.

16. Ayers M, Lunceford J, Nebozhyn M, et al. IFN-gamma-related mRNA profile predicts clinical response to PD-1 blockade. J Clin Invest 2017;127:2930–2940.

17. Cristescu R, Mogg R, Ayers M, et al. Pan-tumor genomic biomarkers for PD-1 checkpoint blockade-based immunotherapy. Science 2018;362.

18. Fehrenbacher L, Spira A, Ballinger M, et al. Atezolizumab versus docetaxel for patients with previously treated non-small-cell lung cancer (POPLAR): a multicentre, open-label, phase 2 randomised controlled trial. Lancet 2016;387:1837–46.

19. Karachaliou N, Gonzalez-Cao M, Crespo G, et al. Interferon gamma, an important marker of response to immune checkpoint blockade in non-small cell lung cancer and melanoma patients. Ther Adv Med Oncol 2018;10:1758834017749748.

20. Vijayan D, Young A, Teng MWL, et al. Targeting immunosuppressive adenosine in cancer. Nat Rev Cancer 2017;17:709–724.

21. Horenstein AL, Chillemi A, Quarona V, et al. NAD(+)-Metabolizing Ectoenzymes in Remodeling Tumor-Host Interactions: The Human Myeloma Model. Cells 2015;4:520–37.

22. Vaisitti T, Audrito V, Serra S, et al. NAD+-metabolizing ecto-enzymes shape tumor-host interactions: the chronic lymphocytic leukemia model. FEBS Lett 2011;585:1514–20.

23. Ohta A. A Metabolic Immune Checkpoint: Adenosine in Tumor Microenvironment. Front Immunol 2016;7:109.

24. Stagg J, Smyth MJ. Extracellular adenosine triphosphate and adenosine in cancer. Oncogene 2010;29:5346–58.

25. Ma SR, Deng WW, Liu JF, et al. Blockade of adenosine A2A receptor enhances CD8(+) T cells response and decreases regulatory T cells in head and neck squamous cell carcinoma. Mol Cancer 2017;16:99.

26. Hatfield SM, Kjaergaard J, Lukashev D, et al. Immunological mechanisms of the antitumor effects of supplemental oxygenation. Sci Transl Med 2015;7:277ra30.

27. Beavis PA, Divisekera U, Paget C, et al. Blockade of A2A receptors potently suppresses the metastasis of CD73+ tumors. Proc Natl Acad Sci U S A 2013;110:14711–6.

28. Mittal D, Young A, Stannard K, et al. Antimetastatic effects of blocking PD-1 and the adenosine A2A receptor. Cancer Res 2014;74:3652–8.

29. Waickman AT, Alme A, Senaldi L, et al. Enhancement of tumor immunotherapy by deletion of the A2A adenosine receptor. Cancer Immunol Immunother 2012;61:917–26.

30. Malavasi F, Deaglio S, Funaro A, et al. Evolution and function of the ADP ribosyl cyclase/CD38 gene family in physiology and pathology. Physiol Rev 2008;88:841–86.

31. Garnelo M, Tan A, Her Z, et al. Interaction between tumour-infiltrating B cells and T cells controls the progression of hepatocellular carcinoma. Gut 2017;66:342–351.

32. Llovet JM, Bru C, Bruix J. Prognosis of hepatocellular carcinoma: the BCLC staging classification. Semin Liver Dis 1999;19:329–38.

33. Henderson JM, Sherman M, Tavill A, et al. AHPBA/AJCC consensus conference on staging of hepatocellular carcinoma: consensus statement. HPB (Oxford) 2003;5:243–50.

34. Edmondson HA, Steiner PE. Primary carcinoma of the liver: a study of 100 cases among 48,900 necropsies. Cancer 1954;7:462–503.

35. Chew V, Chen J, Lee D, et al. Chemokine-driven lymphocyte infiltration: an early intratumoural event determining long-term survival in resectable hepatocellular carcinoma. Gut 2012;61:427–38.

36. Stack EC, Wang C, Roman KA, et al. Multiplexed immunohistochemistry, imaging, and quantitation: a review, with an assessment of Tyramide signal amplification, multispectral imaging and multiplex analysis. Methods 2014;70:46–58.

37. Abel EJ, Bauman TM, Weiker M, et al. Analysis and validation of tissue biomarkers for renal cell carcinoma using automated high-throughput evaluation of protein expression. Hum Pathol 2014;45:1092–9.

38. Lovisa S, LeBleu VS, Tampe B, et al. Epithelial-to-mesenchymal transition induces cell cycle arrest and parenchymal damage in renal fibrosis. Nat Med 2015;21:998–1009.

39. Garnelo M, Tan A, Her Z, et al. Interaction between tumour-infiltrating B cells and T cells controls the progression of hepatocellular carcinoma. Gut 2015;15:2015–310814.

40. Yeong J, Thike AA, Lim JC, et al. Higher densities of Foxp3(+) regulatory T cells are associated with better prognosis in triple-negative breast cancer. Breast Cancer Research and Treatment 2017;163:21–35.

41. Lim JCT, Yeong, J. P. S., Lim, C. J., Ong, C. C. H., Chew, V. S. P., Ahmed, S. S., Tan, P. H., & Iqbal, J.. An automated staining protocol for 7-colour immunofluorescence of human tissue sections for diagnostic and prognostic use. Journal of The Royal College of Pathologists of Australasia In Press.

42. Esbona K, Inman D, Saha S, et al. COX-2 modulates mammary tumor progression in response to collagen density. Breast Cancer Research 2016;18:35.

43. Mlecnik B, Bindea G, Kirilovsky A, et al. The tumor microenvironment and Immunoscore are critical determinants of dissemination to distant metastasis. Science Translational Medicine 2016;8.

44. Nghiem PT, Bhatia S, Lipson EJ, et al. PD-1 Blockade with Pembrolizumab in Advanced Merkel-Cell Carcinoma. New England Journal of Medicine 2016;374:2542–2552.

45. Feng Z, Jensen SM, Messenheimer DJ, et al. Multispectral Imaging of T and B Cells in Murine Spleen and Tumor. The Journal of Immunology 2016;196:3943–3950.

46. Yeong J, Lim JCT, Lee B, et al. High Densities of Tumor-Associated Plasma Cells Predict Improved Prognosis in Triple Negative Breast Cancer. Frontiers in Immunology 2018;9.

47. Mazzaschi G, Madeddu D, Falco A, et al. Low PD-1 Expression in Cytotoxic CD8(+) Tumor-Infiltrating Lymphocytes Confers an Immune-Privileged Tissue Microenvironment in NSCLC with a Prognostic and Predictive Value. Clin Cancer Res 2018;24:407–419.

48. Yeong J, Lim JCT, Lee B, et al. Prognostic value of CD8 + PD-1+ immune infiltrates and PDCD1 gene expression in triple negative breast cancer. J Immunother Cancer 2019;7:34.

49. Fiore C, Bailey D, Conlon N, et al. Utility of multispectral imaging in automated quantitative scoring of immunohistochemistry. Journal of Clinical Pathology 2012;65:496–502.

50. Abel EJ, Bauman TM, Weiker M, et al. Analysis and validation of tissue biomarkers for renal cell carcinoma using automated high-throughput evaluation of protein expression. Human Pathology 2014;45:1092–9.

51. Feng Z, Bethmann D, Kappler M, et al. Multiparametric immune profiling in HPV– oral squamous cell cancer. JCI Insight 2017;2.

52. RStudio: integrated development environment for R. Boston: RStudio, Inc, 2015.

53. R: a language and environment for statistical computing. Vienna: R Foundation for Statistical Computing, 2016.

54. Eisenhauer EA, Therasse P, Bogaerts J, et al. New response evaluation criteria in solid tumours: revised RECIST guideline (version 1.1). Eur J Cancer 2009;45:228–47.

55. Apolo AB, Infante JR, Balmanoukian A, et al. Avelumab, an Anti-Programmed Death-Ligand 1 Antibody, In Patients With Refractory Metastatic Urothelial Carcinoma: Results From a Multicenter, Phase Ib Study. J Clin Oncol 2017;35:2117–2124.

56. Chow LQM, Haddad R, Gupta S, et al. Antitumor Activity of Pembrolizumab in Biomarker-Unselected Patients With Recurrent and/or Metastatic Head and Neck Squamous Cell Carcinoma: Results From the Phase Ib KEYNOTE-012 Expansion Cohort. J Clin Oncol 2016;34:3838–3845.

57. Sul J, Blumenthal GM, Jiang X, et al. FDA Approval Summary: Pembrolizumab for the Treatment of Patients With Metastatic Non-Small Cell Lung Cancer Whose Tumors Express Programmed Death-Ligand 1. Oncologist 2016;21:643–50.

58. Pai-Scherf L, Blumenthal GM, Li H, et al. FDA Approval Summary: Pembrolizumab for Treatment of Metastatic Non-Small Cell Lung Cancer: First-Line Therapy and Beyond. Oncologist 2017;22:1392–1399.

59. Ikeda S, Okamoto T, Okano S, et al. PD-L1 Is Upregulated by Simultaneous Amplification of the PD-L1 and JAK2 Genes in Non-Small Cell Lung Cancer. J Thorac Oncol 2016;11:62–71.

60. Larkin J, Chiarion-Sileni V, Gonzalez R, et al. Combined Nivolumab and Ipilimumab or Monotherapy in Untreated Melanoma. N Engl J Med 2015;373:23–34.

61. Morandi F, Horenstein AL, Rizzo R, et al. The Role of Extracellular Adenosine Generation in the Development of Autoimmune Diseases. Mediators Inflamm 2018;2018:7019398.

62. Chen L, Diao L, Yang Y, et al. CD38-Mediated Immunosuppression as a Mechanism of Tumor Cell Escape from PD-1/PD-L1 Blockade. Cancer Discov 2018;8:1156–1175.

63. Kang J, Park KH, Kim JJ, et al. The role of CD38 in Fcgamma receptor (FcgammaR)-mediated phagocytosis in murine macrophages. J Biol Chem 2012;287:14502–14.

64. Lischke T, Heesch K, Schumacher V, et al. CD38 controls the innate immune response against Listeria monocytogenes. Infect Immun 2013;81:4091–9.

65. Lee HC. Cyclic ADP-ribose and nicotinic acid adenine dinucleotide phosphate (NAADP) as messengers for calcium mobilization. J Biol Chem 2012;287:31633–40.

66. Amici SA, Young NA, Narvaez-Miranda J, et al. CD38 Is Robustly Induced in Human Macrophages and Monocytes in Inflammatory Conditions. Front Immunol 2018;9:1593.

67. Lund FE, Cockayne DA, Randall TD, et al. CD38: a new paradigm in lymphocyte activation and signal transduction. Immunol Rev 1998;161:79–93.

68. Xu L, Chen D, Lu C, et al. Advanced Lung Cancer Is Associated with Decreased Expression of Perforin, CD95, CD38 by Circulating CD3+CD8+ TLymphocytes. Ann Clin Lab Sci 2015;45:528–32.

69. ClinicalTrials.gov. A Study of the Safety and Efficacy of Atezolizumab Administered in Combination with Bevacizumab and/or Other Treatments in Participants With Solid Tumors. Volume 2019, 2016.

70. Quarona V, Zaccarello G, Chillemi A, et al. CD38 and CD157: a long journey from activation markers to multifunctional molecules. Cytometry B Clin Cytom 2013;84:207–17.

