## Supplementary Table 1 for "CD38 is a good predictor of anti-PD-1 immunotherapy responsiveness in hepatocellular carcinoma"

***Supplementary Material***

**Supplementary Tables**

**Supplementary Table 1** List of antibodies used for multiplex immunofluorescence and IHC staining

| **Antibody** | **Source** | **Labeling pattern** |
| --- | --- | --- |
| DAPI | Perkin Elmer (FP1490) | Cell nucleus |
| CD38 | Novo Castra (NCL-CD38-290) | Immune cells, cell membrane |
| CD68 | Dako (M0876) | Immune cells, cytoplasm |
| CD8 | Novo Castra  (NCL-CD8-4B11) | Immune cells, cell membrane |
